# Concomitant effects of orchiectomy and intermittent hypoxia on hepatic oxidative stress, expression of flavin-containing monooxygenases and transcriptomic profile in mice

**DOI:** 10.1101/2023.05.24.541054

**Authors:** Gauthier Ganouna-Cohen, François Marcouiller, Charles Joly Beauparlant, Arnaud Droit, Elise Belaidi, Aida Bairam, Vincent Joseph

**Affiliations:** Centre de Recherche de l Institut Universitaire de Cardiologie et de Pneumologie de Quebec, Universite Laval. Departement de Pediatrie, Faculte de Medecine. Quebec, QC, Canada; Centre de Recherche du Centre Hospitalo-Universitaire de Québec. Département de Médecine Moléculaire, Faculté de Médecine. Québec, QC, Canada; Université Grenoble Alpes, HP2, INSERM, U1300, F-38042, Grenoble, France; Institut de Biologie et Chimie des Protéines UMR5305-LBTI, CNRS, Lyon, France

**Keywords:** Intermittent hypoxia, monooxygenase, FMO3, Liver, Testosterone.

## Abstract

Intermittent hypoxia induces oxidative stress and alters hepatic metabolism, likely underlying the association of sleep apnea with non-alcoholic fatty liver disease. In male patients with sleep apnea, metabolic or liver diseases, the levels of testosterone are reduced, and in patients with metabolic diseases, low levels of testosterone are associated with oxidative stress. To assess potential interactions between testosterone and IH on hepatic oxidative stress we used sham-operated or orchiectomized (ORX) mice exposed to normoxia (Nx) or IH (6% O_2_, 12 cycles/h, 12h/day) for 2 weeks. The activity of prooxidant (NADPH oxidase – NOX), antioxidant enzymes (superoxide dismutase, catalase, and glutathione peroxidase – SOD, Cat, GPx), lipid peroxidation (MDA concentration) and the total concentration of glutathione (GSH) were measured in liver. IH induced a prooxidant profile of enzyme activity (lower SOD activity and higher NOX/SOD and NOX/Cat activity ratio) without altering hepatic MDA and GSH content. Using RNA sequencing followed by a pathway enrichment analysis we identified putative hepatic genes underlying the interactions between IH and testosterone. ORX and IH altered the expression of genes involved in oxidoreductase activities, cytochromes dependent pathways, and glutathione metabolism. Among the genes upregulated in ORX-IH mice, the flavin-containing monooxygenases (FMO) are particularly relevant since these are potent hepatic antioxidant that could help prevent overt oxidative stress in ORX-IH mice.

**Graphical Abstract:** 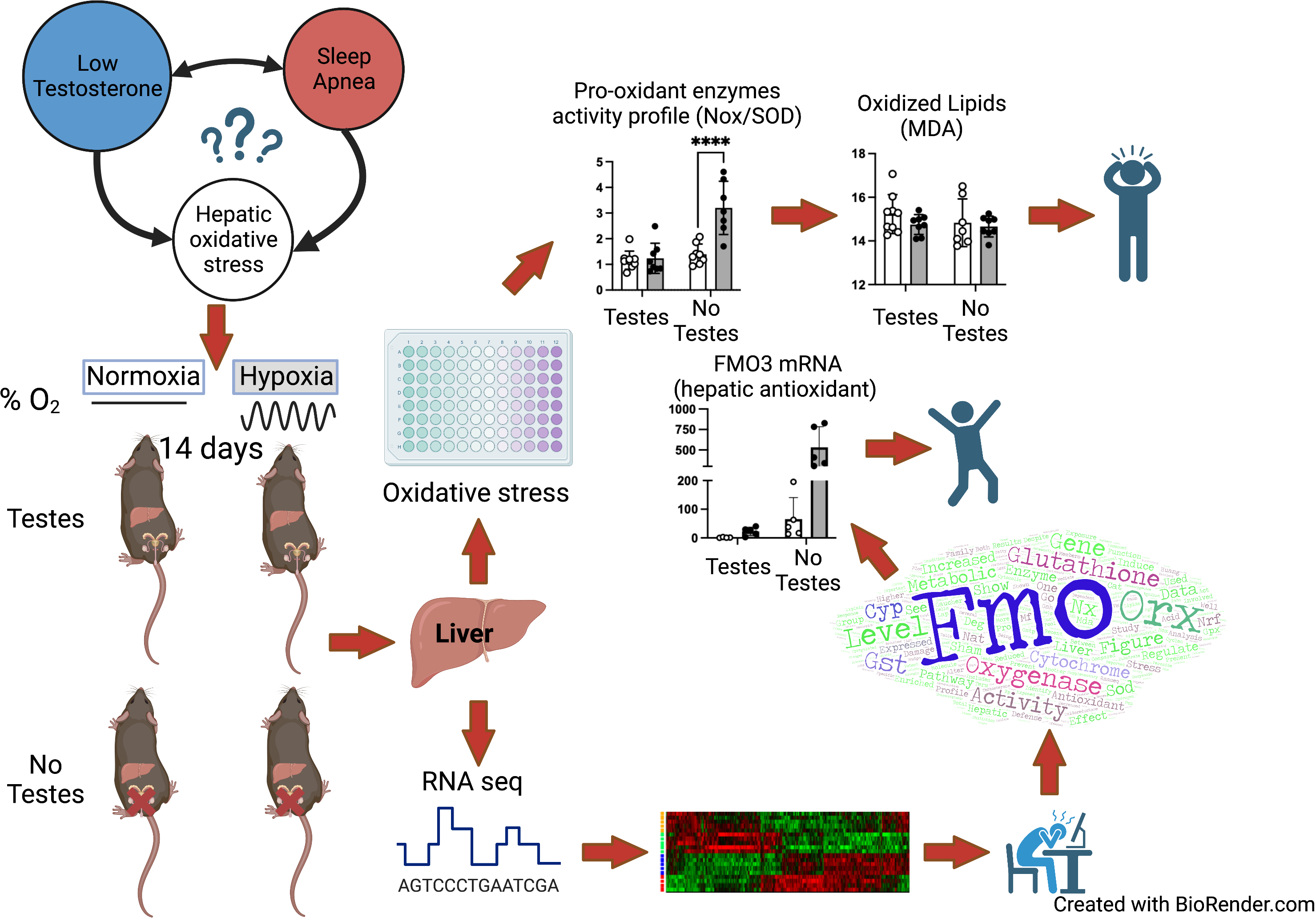

## INTRODUCTION

Sleep apnea (SA) is a highly prevalent respiratory disorder associated with metabolic dysfunctions (Heinzer et al., 2018; Hirotsu et al., 2019) and non-alcoholic fatty liver disease (J. Huang et al., 2023). The association between SA and metabolic diseases is clearly attributable to the episodes of nocturnal intermittent hypoxia (IH) that increase the activity of the sympathetic nervous system and induce insulin resistance (Conde et al., 2023; Ryan, 2017) as well as oxidative stress in the liver (Gaucher et al., 2022; Jun et al., 2008) and adipose tissue (Gileles-Hillel et al., 2017). In the liver, oxidative stress induced by IH occurs despite an increased expression of superoxide dismutase (SOD - one of the main antioxidant enzymes) and specific enrichment of genes regulated by the Nuclear Respiratory Factor 1 and 2 (NRF1 – NRF2, two of the main transcription factors that orchestrate cellular and mitochondrial responses to oxidative stress) as assessed by transcriptomic analysis (Gaucher et al., 2022).

Age is an important risk factor for SA (Heinzer et al., 2015). It is noteworthy that age is associated with reduced level of testosterone and a significant proportion of aging men have extremely low testosterone levels (Kelsey et al., 2014). Coincidently, recent studies have shown that the incidence of hypogonadism is elevated in male SA patients (Kim & Cho, 2019) and it is also acknowledged that low levels of testosterone are related to the occurrence of metabolic hepatic diseases (Yim et al., 2018) and type II diabetes (Rovira-Llopis et al., 2017). In men, testosterone is involved in the regulation of several metabolic processes and improves antioxidant defense mechanism, as illustrated by a negative correlation between blood testosterone concentration and mitochondrial production of reactive oxygen species on one hand and, on the other hand, a positive correlation between testosterone and the mRNA expression level of superoxyde dismutase in leukocytes of type 2 diabetic patients (Rovira-Llopis et al., 2017).

This raises the intriguing hypothesis that testosterone contributes to the regulation of oxidative stress and hepatic metabolic functions in SA patients. To assess this hypothesis, we used intact and orchiectomized (ORX) male mice exposed to control conditions (normoxia: Nx) or IH (the main feature of OSA) for 2 weeks.

We then used liver samples to measure the activities of pro-(NADPH oxidase) and antioxidant enzymes (SOD and catalase - Cat) and assess lipid peroxidation. Because Glutathione (GSH) is one of the most prevalent antioxidant system in the liver that is altered in hepatic diseases (Santacroce et al., 2023), we also assessed the total concentration of GSH as well as the activity of glutathione S-transferase (GST) and glutathione peroxidase (GPx). We then performed unbiased RNA sequencing and a pathway enrichment analysis to uncover potential specific interactions between IH and ORX at a genome-wide level of mRNA expression.

Our results show that in ORX mice, IH induces a pro-oxidant profile of enzyme activity without increasing the concentration of MDA. The RNAseq analysis shows altered expression of a series of genes involved in glutathione metabolism, cytochrome P450 and redox pathways in response to IH, ORX or combined exposures in ORX-IH mice. A particularly interesting finding is that the potent hepatic antioxidants flavin monoamine oxidases (FMO) 1 to 4 are upregulated and might thus provide additional defense mechanisms against oxidative stress specifically in ORX-IH mice.

## MATERIALS AND METHODS

### Animals and ethical approval

All experiments have been approved by the animal protection committee of Université Laval in accordance with the Canadian Council on Animal Care in Science (project # 2019-368). We used a total of 32 C57BL/6NCrl male mice ordered from Charles-River Laboratories (Saint-Constant, QC, Canada) at 4 weeks of age. All mice had *ad-libitum* access to food and water and were maintained on a 12/12h light/dark cycle. All mice were acclimatized to their new environment during four weeks after their arrival to our animal house facility and randomly assigned to the sham operation (Sham) or orchiectomy (ORX) surgery.

### Bilateral orchiectomy

All surgery were performed as previously described (Ganouna-Cohen et al., 2022, 2023). Each mice received a dorsal injection of meloxicam (1 mg/kg), buprenorphine (0.05 mg/kg), and Ringer’s lactate (10 ml/kg/h) before being anesthetised with isoflurane (4% for induction, 2% thereafter, in 30% O_2_). The level of anaesthesia was assessed by the lack of reflex responses to pinches of toes. The scrotum was shaved and opened after injecting 0.05 ml of local anaesthesia (0.875 mg/ml lidocaine + 0.437 5mg/ml bupivacaine). Testes were externalized and removed (for ORX) or returned to the scrotum for the Sham operation. The scrotum was then closed with surgical clips. All mice received a dorsal injection of meloxicam (1 mg/kg) the day after the surgery and were subjected to close daily observation for 5 additional days. Surgical clips were removed 7 days after the surgery.

### Intermittent hypoxia

Two weeks following the surgery (i.e. 10 weeks old), mice were housed in standard cages (4 mice/cage) connected to an oxycycler (Biospherix, Redfield, NY, USA). Oxygen dropped from 21% to 6% in 90 seconds, remained at 6% for 30 seconds, then returned to 21% in 70 seconds, and remained at normoxia for 120 seconds. There were 12 cycles/hour for 12 consecutive hours between 6:00 am - 6:00 pm for 14 consecutive days, as described previously (Ganouna-Cohen et al., 2022, 2023; Marcouiller et al., 2021). Following the last day of exposure to IH or Nx, the mice were fasted overnight for 12 hours (9 pm – 9 am) then allowed to feed ad-libitum for 2 hours. We used this strategy to ensure as much as possible similar food ingestion across groups before evaluation of liver oxidative stress and gene expression profile. The mice were then weighed and euthanized with an overdose of anesthetic (ketamine/xylazine) followed by a cardiac puncture, the liver was rapidly dissected, immediately frozen and kept at -80°C until later use.

### Measurements of Oxidative Stress Enzyme Activities

We used commercially available kits to measure the concentration of malondialdehyde (MDA – TBARS assay, R&D assays, cat #KGE013), the total concentration of glutathione (GSH – Cayman chemicals, cat #703002) and the activity of glutathione S-transferase (Cayman Chemical, cat #703302). All kits were used as described by the recommended protocols for extraction and assay of soft tissue, and values were expressed relative to wet liver mass.

We measured the activity of NADPH oxidase (NOX), superoxide dismutase (SOD), catalase (Cat), and glutathione peroxidase (GPx) using standard protocol as previously described (Jochmans-Lemoine et al., 2018; Laouafa et al., 2017; Ribon-Demars et al., 2021) and detailed below.

### Protein Extraction

After homogenization of the frozen liver samples in a phosphate buffer, different centrifugations (all at 4°C) were used to separate cytosolic and mitochondrial fractions. Briefly, the samples were centrifuged for 4 minutes at 1,500 g followed by a second centrifugation for 10 minutes at 12,000g to collect the cytosolic fraction (supernatant). The pellet was then re-suspended in a mitochondrial isolation buffer (250 mM sucrose, 1 mM EGTA, 20 mM Tris base, pH 7.3) and centrifuged for 10 minutes at 1,500g, the supernatant was recovered and centrifuged at 9,000g for 11 minutes. The pellet (mitochondrial fraction) was re-suspended in 200 μl of mitochondrial isolation buffer. These fractions were stored at -80°C until analyses. The concentration of proteins in each fraction was determined by a standard colorimetric BCA assay kit (ThermoFisher Scientific, catalogue # 23225). All assays were performed in duplicate on 96 wells plate with an equal quantity of purified protein into each well (200 µg and 2 µg for cytosolic and mitochondrial fractions respectively) and read on a SpectraMax Microplate reader (Molecular Devices, San Jose, CA, USA).

### NADPH oxidase (NOX)

NOX activity was assessed using a cocktail containing nitroblue tetrazolium (NTB, 2.2 mM in water), Tris–HCL pH 8 (2.8 mM), and diethylene-triamine-penta-acetic acid (1.3 mM in Tris–HCL). The reaction mix was composed of 20 μl of sample, 250 μl of cocktail, and 30 μl (100 μM per well) of a fresh NADPH solution (1mM). The plate was shaken 2 minutes at room temperature, then the absorbance was read at 450 nm every 30 seconds for 10 minutes. NOX activity corresponds to the rate of formation of formazan blue (calculated with the extinction coefficient of 0.019 mM^-1^/cm^-1^) and expressed as nmol/min/µg protein.

### Superoxide dismutase (SOD)

The activity of SOD was measured in the cytosolic and mitochondrial fractions. Cytosolic SOD activity was determined by the degree of inhibition of the reaction between O_2_^・-^ produced by a hypoxanthine/xanthine oxidase system and NTB. We used the cocktail detailed for the NOX activity plus hypoxanthine (0.19 mM). A fresh solution of xanthine oxidase (1.02 units/ mL) was prepared, then we added 20 μl of sample, 250 μL of cocktail, and 20 μl of xanthine oxidase to each well. The plate was mixed for 4–5 seconds at room temperature. The absorbance was quickly read at 450 nm every 30 seconds for 6 minutes. SOD activity was determined in the mitochondrial fraction with 1 mM of NaCN in each well to inhibit the cytosolic SOD. For each assay, four wells were used as blanks, with 20 μl of PBS 1x rather than samples. The rate of appearance of NTB was determined in the blanks and in the samples using the extinction coefficient of 0.019 mM^-1^.cm^-1^. SOD activity corresponded to the difference between these slopes, expressed as nmol/min/µg protein.

### Catalase (Cat)

The activity of Cat was determined by the formation rate of formaldehyde with methanol and H_2_O_2_ as substrates. We added 20 µl of samples, 100 µl PBS 1x, 30 µl methanol (100%) and 20 µl H_2_O_2_ (0.15%) into each well and let the reaction run for 20 minutes. The reaction was stopped with 30 µl KOH 10N, and we added 30 µl of a 0.2 M purpald solution to determine the concentration of formaldehyde at 540 nm against a standard curve (0-300 µM). The activity of Cat was determined as nmol/min/µg protein.

### Glutathione peroxidase (GPx)

The activity of GPx was determined as the rate of oxidation of NADPH to NADP+ in a cocktail solution containing glutathione reductase, NADPH (1.7 mM), and reduced glutathione (1.6 mM) using a solution of cumene hydroperoxide (15 µM) as substrate. We added 20 μl of sample, 200 μl of PBS 1x, 30 μl of the cocktail solution, and 30 μl of cumene hydroperoxide solution to each well and mix the plate 4–5 seconds at room temperature. The absorbance was quickly read at 340 nm every 50 seconds for 5 minutes. GPx activity was measured as the rate of NADPH consumption (using the extinction coefficient of 6.22 mM^-1^.cm^-1^) and expressed as nmol/min/µg protein.

### Transcriptomic analysis of liver mRNA expression. Extraction of RNA, cDNA synthesis and sequencing

We used 5 mice from each group for total RNA extraction from frozen liver samples using EasyPure® RNA Kit (TransGen Biotech Co. Cat. No ER 101 China) according to the manufacturer’s instructions. Briefly, frozen samples were weighed and mechanically homogenized in binding buffer (300 µl/10 mg), a proteinase K solution (15µl/10 mg) and β-mercaptoethanol (1%), incubated for 15 minutes at 56°C and centrifuged at 12,000g 5 min at room temperature. Then RNA was purified by adding an equal volume of 70% ethanol to the lysate and vortexed. It was then centrifuged 20 sec at 12,000g, the lysate was added into the spin columns and re-centrifuged 30 sec at 12,000g. The columns were cleaned (500 µl of clean buffer, centrifuged at 12,000g for 35 sec), digestion of DNA was performed by adding 80 µl of DNAse I for 15 minutes, cleaned, and washed twice (500 µl of washing buffer, centrifuged at 12,000g for 35 sec), followed by a centrifugation at maximum speed 2 min. The column was then air-dried 3 min and RNA was eluted with water (100 µl). Total RNA concentration was determined by the absorption at 260 nm using a Thermo Scientific^TM^ NanoDrop^TM^ 2000c spectrophotometer (Thermo Fisher Scientific, MA, USA). The 260/280 nm and 230/260 nm absorption ratio were assessed. All samples had absorption ratios of 260/280 > 1.8 and 260/230 typically >1.3. RNAs were then stored at -80°C until being used for RNA sequencing.

### RNA sequencing

All procedures have been carried out by Génome Québec for the RNA sequencing according to their standard protocols. Briefly, total RNA was quantified, and its integrity was assessed using 5K / RNA / Charge Variant Assay LabChip and RNA Assay Reagent Kit (Perkin Elmer). Libraries were generated from 250 ng of total RNA as following: mRNA enrichment was performed using the NEBNext Poly(A) Magnetic Isolation Module (New England BioLabs). cDNA synthesis was achieved with the NEBNext RNA First Strand Synthesis and NEBNext Ultra Directional RNA Second Strand Synthesis Modules (New England BioLabs). The remaining steps of library preparation were done using the NEBNext Ultra II DNA Library Prep Kit for Illumina (New England BioLabs). Adapters and PCR primers were purchased from New England BioLabs.

The libraries were normalized, pooled, and then denatured in 0.02N NaOH and neutralized using HT1 buffer. The pool was loaded at 175 pM on an Illumina NovaSeq S4 lane using Xp protocol as per the manufacturer’s recommendations. The run was performed for 2x100 cycles (paired-end mode). A phiX library was used as a control and mixed with libraries at 1% level. Base calling was performed with RTA v3. The bcl2fastq2 v2.20 program was then used to demultiplex samples and generate fastq reads.

Reads were trimmed using fastp v0.23.2 (Chen et al., 2018). Quality check was performed on raw and trimmed data to ensure the quality of the reads using FastQC v0.11.9 (Andrews, 2010) and MultiQC v1.12 (Ewels et al., 2016). The quantification was performed with Kallisto v0.46.1 (Bray et al., 2016) against the Mus musculus transcriptome (Ensembl release 108). Raw reads were analyzed using the Qlucore omic explorer software (Qlucore, Lund, Sweden). The data table was imported in Qlucore and normalized using the trimmed mean of M-values (TMM), a method that takes into account differences of the underlying distribution of expressed transcripts to adequately compare gene expression between different experimental conditions (Robinson & Oshlack, 2010).

### Statistical analyses and data presentation

Enzyme activities were analyzed with 2-way ANOVAs using ORX and IH as grouping variables. If significant effects or interactions appeared, a post-hoc Fisher’s LSD analysis was used to assess significant effects between groups. All values are reported as mean ± SD. All analyses have been done with Graphpad Prism for macOS (version 9.5.0).

For the RNAseq analysis we applied a one-way ANOVA to discriminate genes differentially expressed between groups with a significant threshold of 0.1 corrected for the false discovery rate (q value – equivalent p value = 0.0069). From this set of genes, we generated a heat table and a principal component analysis (PCA). The list of differentially expressed genes (DEG) along with the results of the ANOVA and a Tukey’s range post-hoc test to assess differences between groups was exported to an excel spreadsheet. We used this list of DEG to perform an enrichment analysis with the g:Profiler web server with a g:SCS multiple testing correction method applying a significance threshold of 0.05 (Raudvere et al., 2019). For functional analysis, we selected the Gene Ontology data base for molecular function and biological function without electronic annotation (GO:MF, GO:BF) and the Kyoto Encyclopedia of Genes and Genomes (KEGG). Specific lists of DEG for each relevant comparison (effect of IH in sham mice: Sham IH vs Sham Nx / effect of ORX: ORX Nx vs Sham Nx / effect of IH in ORX mice: ORX IH vs ORX Nx) were then extracted and used to assess the number of genes up or down regulated in each condition and construct a Venn Diagram (with the Qlucore tool) to show potentially overlapping DEG. When comparing expression levels of a given gene between two groups, fold changes were calculated from the TMM reads value as e^(0.6931 x TMMDiff)^, where TMMDiff is the difference of TMM count reads between these groups.

## RESULTS

Body weight and plasma testosterone levels are presented in table 1. As previously reported (Ganouna-Cohen et al., 2022, 2023) ORX successfully reduced testosterone levels, and both IH and ORX reduced body weight (ANOVA p value respectively < 0.0001 and 0.006).

**Table 1:** Body weight and plasma testosterone levels measured in Sham and ORX mice exposed to Nx or IH. All values are mean ± SD. Number of mice in each group indicated as (n=). P value for post-hoc analysis: **, ****: p < 0.01 and <0.0001 IH vs Nx / †, †††: p < 0.05 and <0.001 ORX vs Sham

|  | Sham Nx<br>(n=9) | Sham IH<br>(n=8) | ORX Nx<br>(n=8) | ORX IH<br>(n=7) |
| --- | --- | --- | --- | --- |
| Body weight<br>(g) | 26.2 ± 1.4 | 22.2 ± 1.3<br>**** | 23.5 ± 1.8<br>††† | 21.5 ± 0.9<br>** |
| Testosterone<br>(ng/ml) | 4.91 ± 3.36 | 2.72 ± 3.23 | 0.08 ± 0.08<br>††† | 0.06 ± 0.03<br>† |

### IH exposures induce a cytosolic and mitochondrial pro-oxidant profile of enzymatic activity in ORX mice

In sham mice, the cytosolic SOD activity (cyto. SOD) was increased by IH exposures (P value for IH x ORX = 0.0001), without changing the activity of the other enzymes (Figure 1 A-C). By contrast, in ORX mice, the cyto. SOD activity was reduced by IH, showing lower antioxidant defense. The resulting NOX/SOD and NOX/Cat ratio were about 2-3 times higher in ORX-IH mice compared to the other groups (Figure 1 D & E) showing a bias towards pro-oxidant activities. However, the MDA concentration in ORX-IH mice was comparable to the other groups (Figure 1 F), suggesting that defenses preventing the built-up of oxidized lipids are still efficient. In the mitochondrial fraction of Sham mice, IH reduced CAT activity (p value for IH x ORX = 0.024), without altering SOD or GPx activities (Figure 1 G-I). In ORX mice there was a non-significant trend towards decreased SOD activity induced by IH (p value for IH x ORX = 0.047).

**Figure 1:**
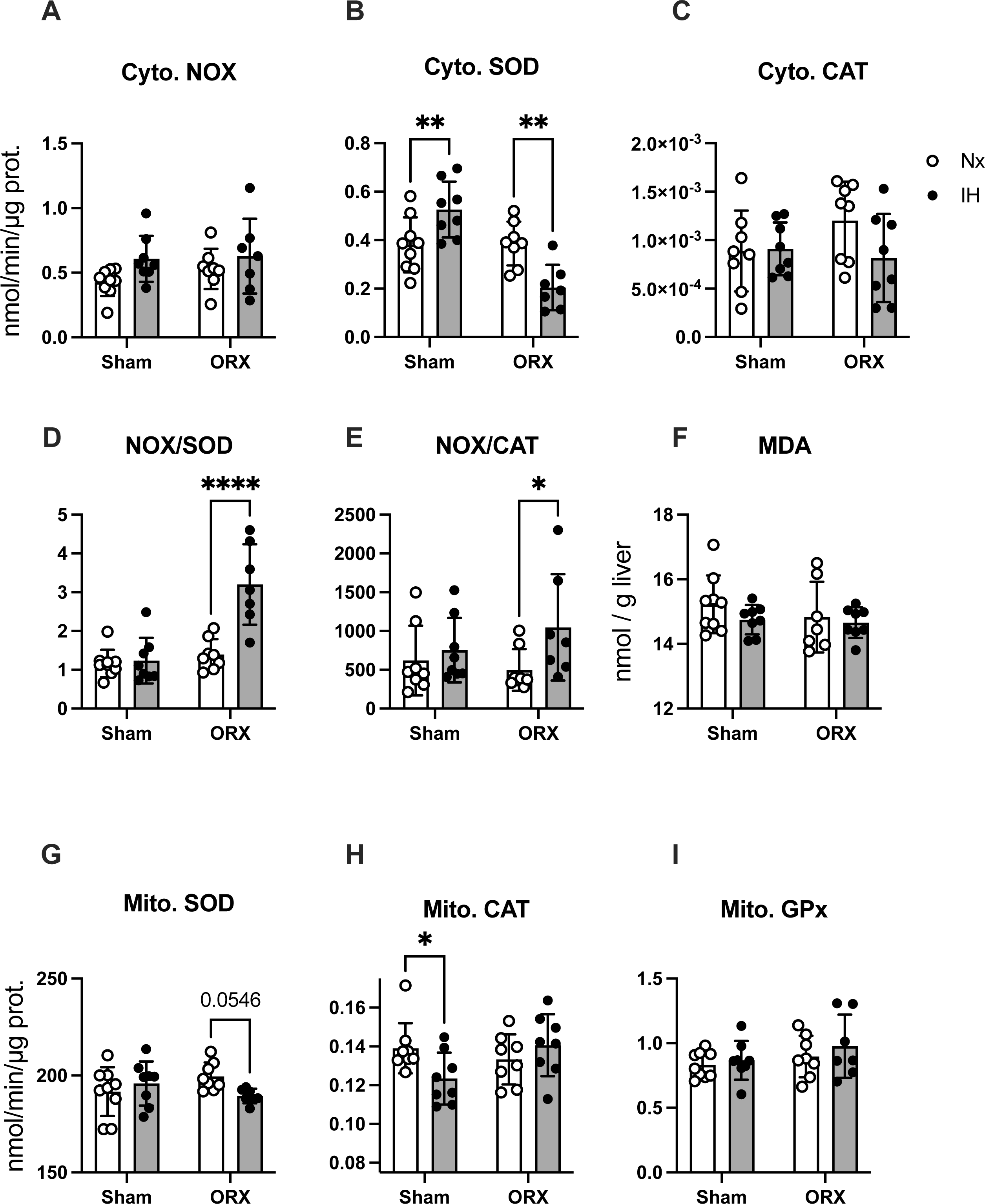
Profile of enzymatic activities in the liver of Sham and ORX mice exposed to Nx or IH. **A-C**: activity of NADPH oxidase (NOX), superoxide dismutase (SOD) and catalase (CAT) in the cytosolic fraction (Cyto. – all in nmol/min/µg prot). **D-E**: ratio of NOX/SOD and NOX/CAT and concentration of MDA (in ng/g liver). **G-I**: Activity of superoxide dismutase (SOD), catalase (CAT) and glutathione peroxidase (GPx) in the mitochondrial fraction (Mito. – all in nmol/min/µg). All values are mean ± SD. Individual data points are shown. P-values for post-hoc analysis: *, **, ****: p<0.05, <0.01, and <0.0001 IH vs Nx.

The total GSH concentration and GPx activity were not altered by IH or ORX (Figure 2), but GST activity was reduced by IH (p = 0.008), with the post-hoc analysis showing significant effect only when comparing ORX-IH vs ORX-Nx mice.

**Figure 2:**
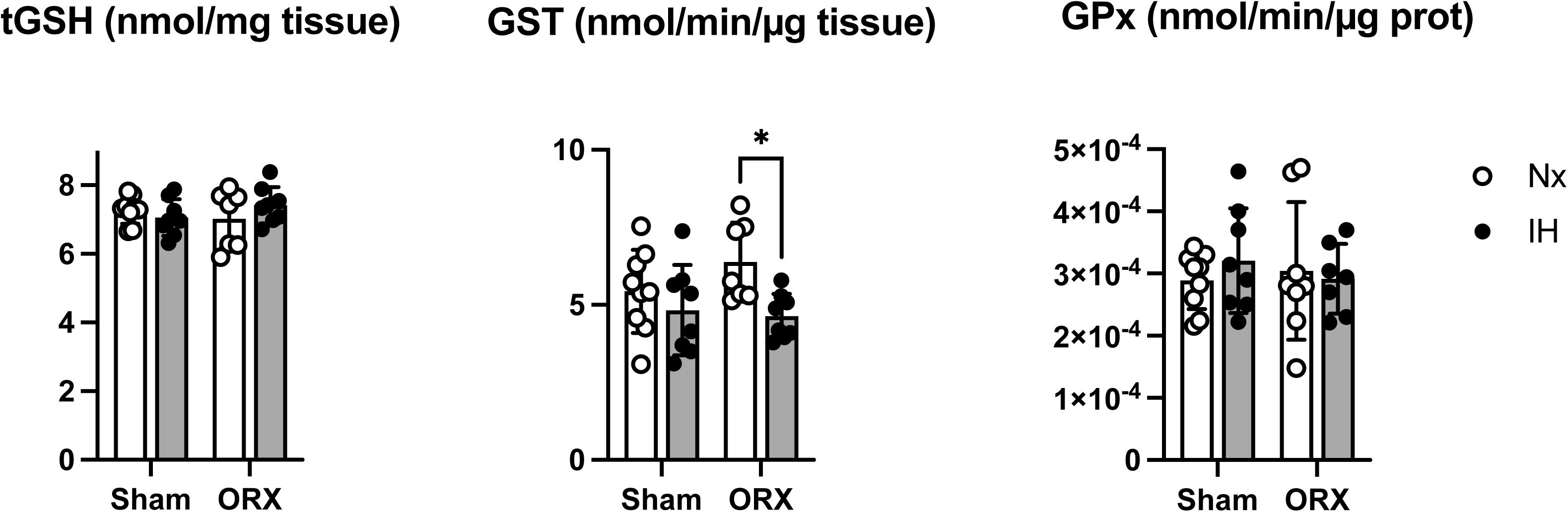
Cytosolic glutathione metabolism in the liver of Sham and ORX mice exposed to Nx or IH. **A:** Total glutathione concentration (tGSH – nmol/mg liver). **B:** Activity of glutathione-S-transferase (GST – nmol/min/µg tissue). **C**: activity of glutathione peroxidase (GPx – nmol/min/µg protein). All values are mean ± SD. Individual data points are shown. P-values for post-hoc analysis: *: p<0.05 IH vs Nx.

### RNAseq analysis reveals alternative antioxidant pathways in ORX-IH mice through regulation of flavin-containing monooxygenase

To explain the particular apparent protection against accumulation of oxidized lipids despite a pro-oxidant profile of enzymatic activity, we used an unbiased approach to evaluate the effects of IH and ORX at the gene expression level by using a RNAseq analysis (with 5 animals/group). One sample from the Sham-Nx group did not pass the quality test and was excluded from further analysis. The gene expression pattern across the different groups is presented in Figure 3. There was a total of 2485 differentially expressed genes (DEG – Figure 3 A – see supplementary data 1 for expression data and statistics) in the liver with most of these genes (about 1700 – 68%) being regulated by ORX in normoxic conditions (Figure 3 B). The PCA plot (Figure 3 C) shows tight clustering of individuals within each experimental group, and the Venn diagram (Figure 3 D) shows that a limited number of genes are commonly regulated by IH in sham and ORX mice (around 25%).

**Figure 3:**
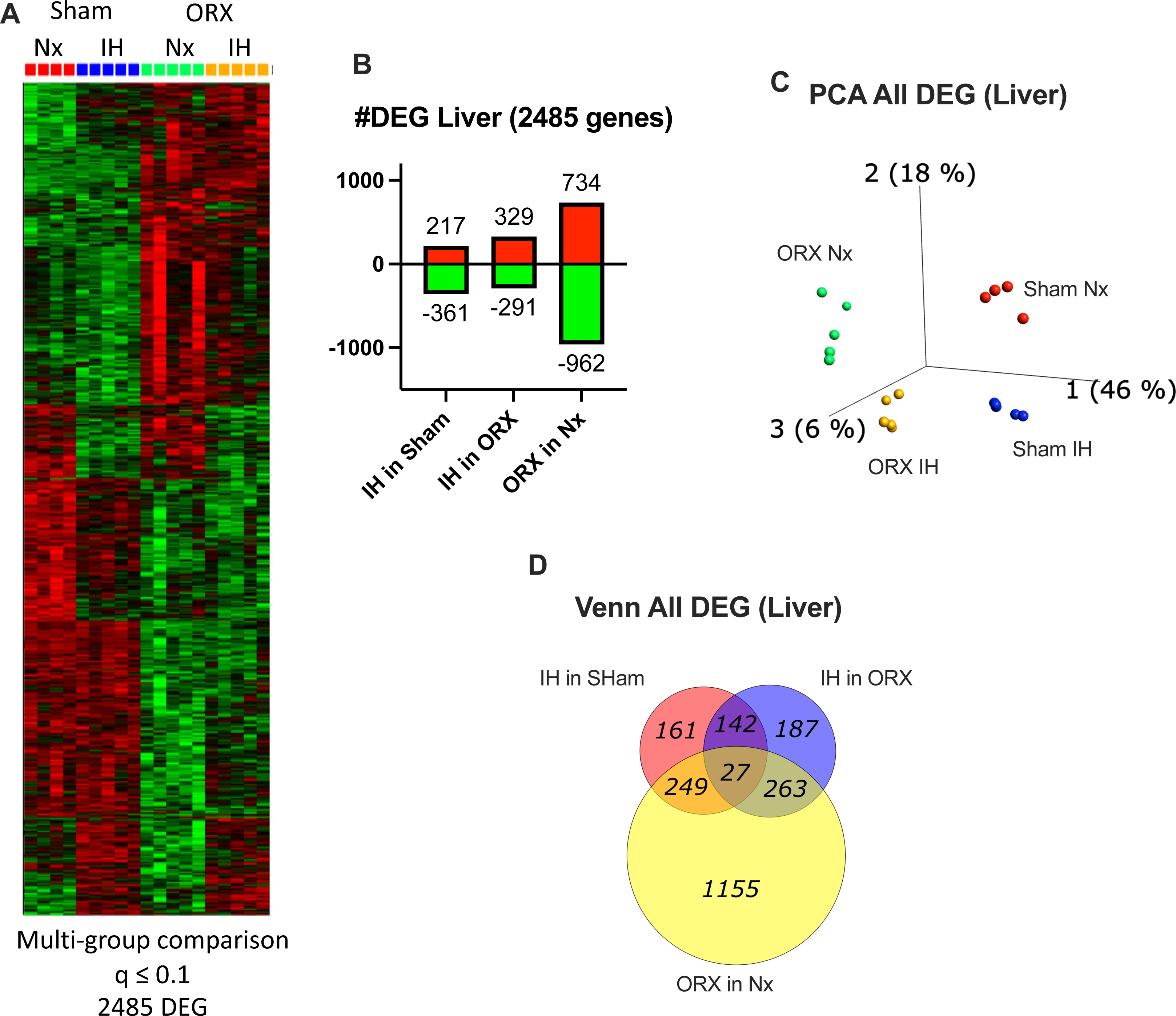
Transcriptomic profile of RNA expression in the liver of Sham and ORX mice exposed to Nx or IH (RNA sequencing) **A:** Heat table and hierarchical clustering of differentially expressed genes (DEG – adjusted p value <0.1 – upregulation in red, down regulation in green). **B:** Number of upregulated (red and positive numbers) and downregulated (green and negative numbers) genes by IH in sham mice (Sham IH vs Sham Nx) or ORX mice (ORX IH vs ORX Nx) and by ORX in Nx mice (ORX Ns vs Sham Nx). **C:** Principal component analysis (CPA – 3D score plots of principal components 1-3) for individual mice showing tight clustering by group. **D:** Venn diagram of DEGs showing number of overlapping genes for the effects of IH in Sham or ORX mice and the effect of ORX in sham mice.

Among the DEG identified, there were highly significant enrichments for oxidoreductase and monooxygenase activities (GO:MF Figure 4 A), a series of metabolic processes (GO:BP Figure 4 B), as well as cytochromes dependent pathways and glutathione metabolism (KEGG - Figure 4 C – see supplementary data 2 for the complete list of enriched pathways). These pathways appear particularly relevant given the effects of IH and ORX on the profile of enzymatic activities involved in oxidative stress, we therefore sought to identify more precisely how these changes are related to the oxidative stress pathways.

**Figure 4:**
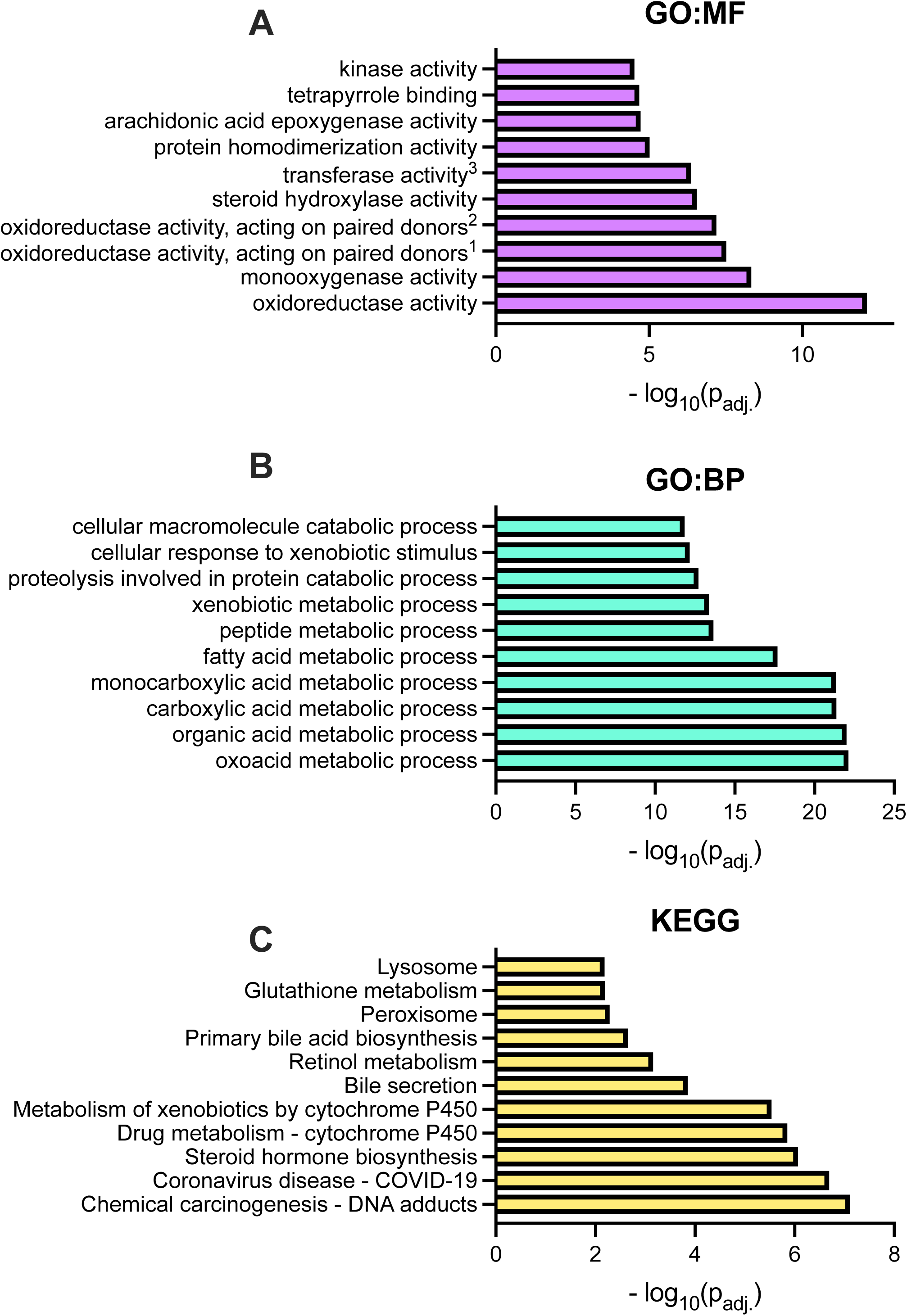
Enrichment analysis of DEG identified in the liver of Sham and ORX mice exposed to Nx or IH with the GO:MF. (**A**), GO:BP (**B**) and KEGG (**C**) annotations. For each graph we selected the top 10 enriched pathways. X-axis are -log_10_ of the adjusted P values for each pathway. Complete definition of terms in GO:MF as follows: ^1^: oxidoreductase activity, acting on paired donors, with incorporation or reduction of molecular oxygen ^2^: oxidoreductase activity, acting on paired donors, with incorporation or reduction of molecular oxygen, reduced flavin or flavoprotein as one donor, and incorporation of one atom of oxygen ^3^: transferase activity, transferring phosphorus-containing groups

A subset of 22 DEGs are involved in glutathione metabolism (see supplementary data 3 for expression data and statistics). Several members of the GPx and GST families were affected either by IH or ORX, including GPX4, the GST π1, π2 (Gstp1 and Gstp2) and the microsomal GST1 (Mgst1) that are downregulated by ORX. On the other hand, GST θ3 (Gstt3) and GST µ7 (Gstm7) are upregulated by ORX (Figure 5 A and B), while GPX7 is upregulated by ORX but reduced by IH in the ORX mice. It is also noteworthy that the expression levels of 6 phosphogluconate dehydrogenase (Pgd) and the mitochondrial isocitrate dehydrogenase 2 (Idh2) are increased by ORX (Figure 5 C): these enzymes are involved in the reduction of oxidized glutathione (GSSG) to renew the GSH pool by providing NADPH molecules that are necessary for the activity of Glutathione reductase (however, the expression level of Glutathione reductase was not altered by IH or ORX – Figure 5 C).

**Figure 5:**
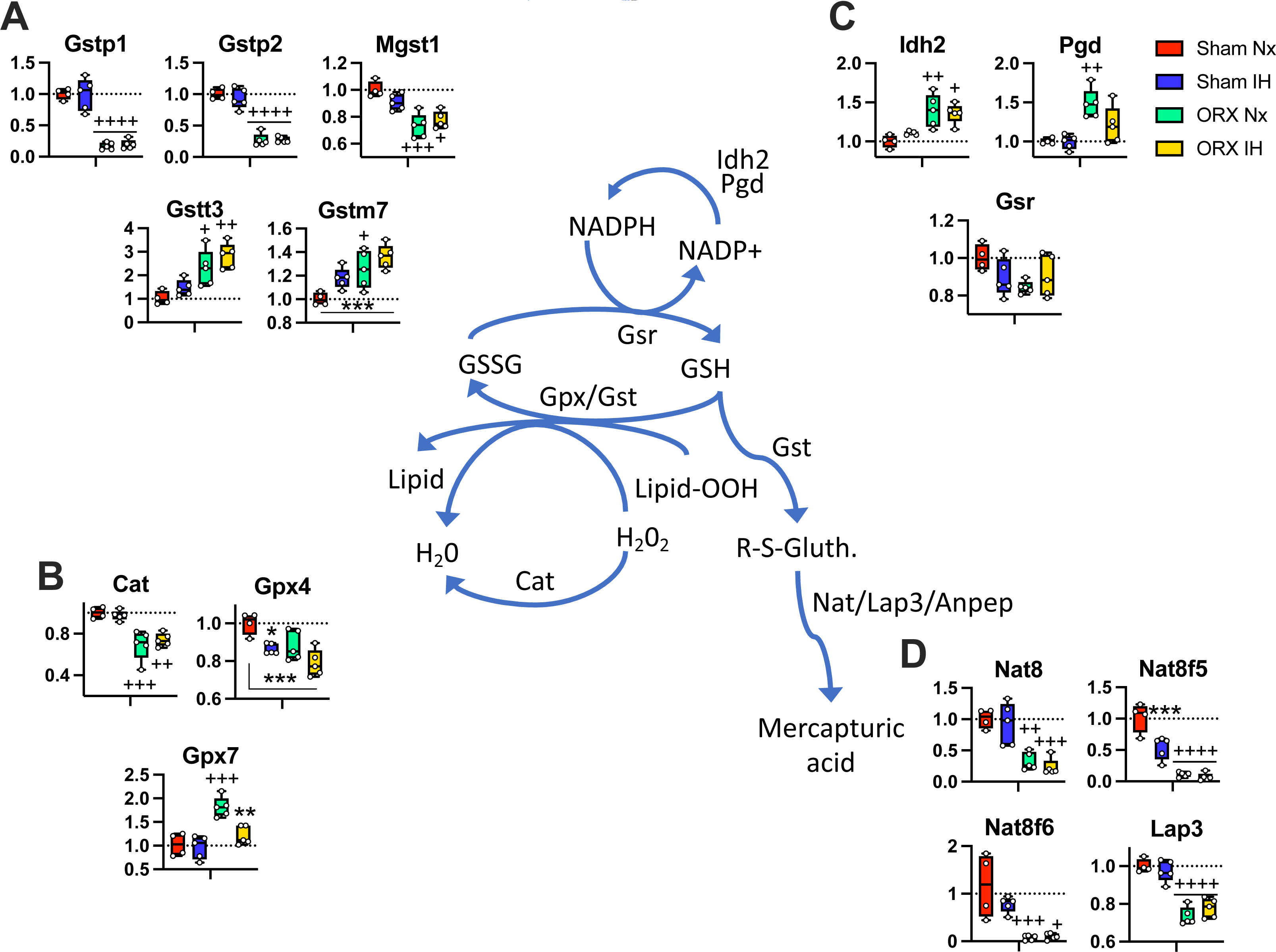
Representative pathway of glutathione metabolism showing the relative expression of DEG identified in the liver of Sham and ORX mice exposed to Nx or IH. See text for further details of gene identification and functions. All values are fold change relative to Sham Nx. P-values for post-hoc analysis: *, ***: p<0.05, p<0.001 IH vs Nx. +, ++, +++, ++++: p<0.5, p<0.01, p<0.001, and p<0.0001 ORX vs Sham.

Another important line of defense against reactive electrophiles molecules is mediated by their conjugation with glutathione and excretion as mercapturic acid through the pathway catalyzed by GST and Nat/Lap3/Anpep (Hanna & Anders, 2019; Ntamo et al., 2021). This branch of mercapturic acid synthesis was also affected by IH and ORX, with decreased expression of N-acetyltransferase 8 (Nat8), Nat8 family members 5 and 6 (Nat8f5, Nat8f6) and leucine aminopeptidase 3 (Lap3 - Figure 5 D) in ORX mice in addition to the altered expression levels of GST family members described above. Finally, it is also noteworthy that the expression level of catalase (Cat) that reduces hydrogen peroxide to water independently of GSH metabolism was also reduced by ORX.

We then extracted the DEG from the GO:MF terms related to oxidoreductase or monooxygenase activity (see supplementary data 4 for expression data and statistics). Among these DEG (representing 140 genes), the most strongly regulated genes were cytochromes responding to orchidectomy (Cyp2b13, Cyp2b9, Cyp3a44 and Cyp2a4 upregulated - Cyp4a12b and Cyp4a12a downregulated – Figure 6 A-C) that oxidize steroids, fatty acids, and xenobiotics and are playing important roles in liver metabolic and detoxification functions. Among the genes with the highest upregulation in ORX-IH mice, the liver antioxidant FMO3 (flavin-containing monoamine oxidase 3) is particularly intriguing: compared to Sham-NX mice, FMO3 expression levels were respectively 23, 65 and 532 times higher in Sham-IH, ORX-NX and ORX-IH mice (Figure 6 A and B). Since FMO3 belongs to a family of enzymes that have been reported as important antioxidant (S. Huang et al., 2021), we assessed whether other FMOs were expressed or regulated in our samples. To answer this, we used the unfiltered data table (containing all the reads aligned to known genes in the RNAseq library) and identified 6 others FMO enzymes, among which 4 were expressed at moderate or high levels (FMO1, FMO2, FMO4, and FMO5), while FMO 6 and 9 remained at very low levels across groups (Table 2). Fold changes relative to Sham-Nx mice for FMO 1-5 are presented in Figure 7, showing higher expression of FMO 1, 2, 3 and 4 in ORX-IH compared to Sham-Nx mice, and higher expression of FMO 2, 3 and 4 in ORX-IH compared to ORX-Nx mice.

**Figure 6:**
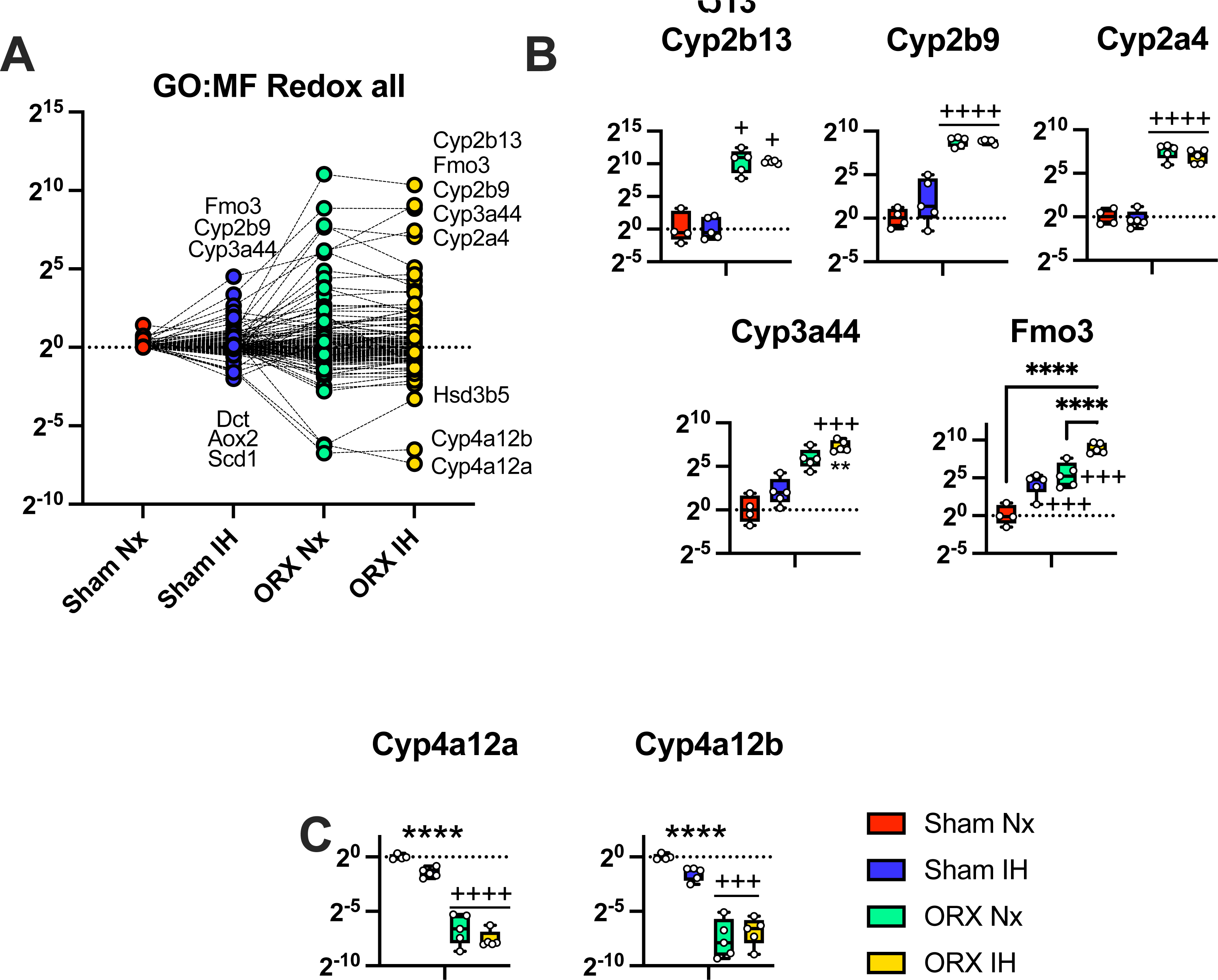
Relative expression of DEG involved in Redox pathways as identified in the GO:MF enrichment analysis in the liver of Sham and ORX mice exposed to Nx or IH. **A:** all genes related to Redox pathways, with few individual names from the most strongly expressed genes. **B and C:** expression pattern of selected genes involved in RedOx pathways. See text for further details of gene identification and functions. All values are fold change relative to Sham Nx. P-values for post-hoc analysis: **, ****: p<0.01, p<0.0001 IH vs Nx. +, +++, ++++: p<0.5, p<0.001, and p<0.0001 ORX vs Sham.

**Figure 7:**
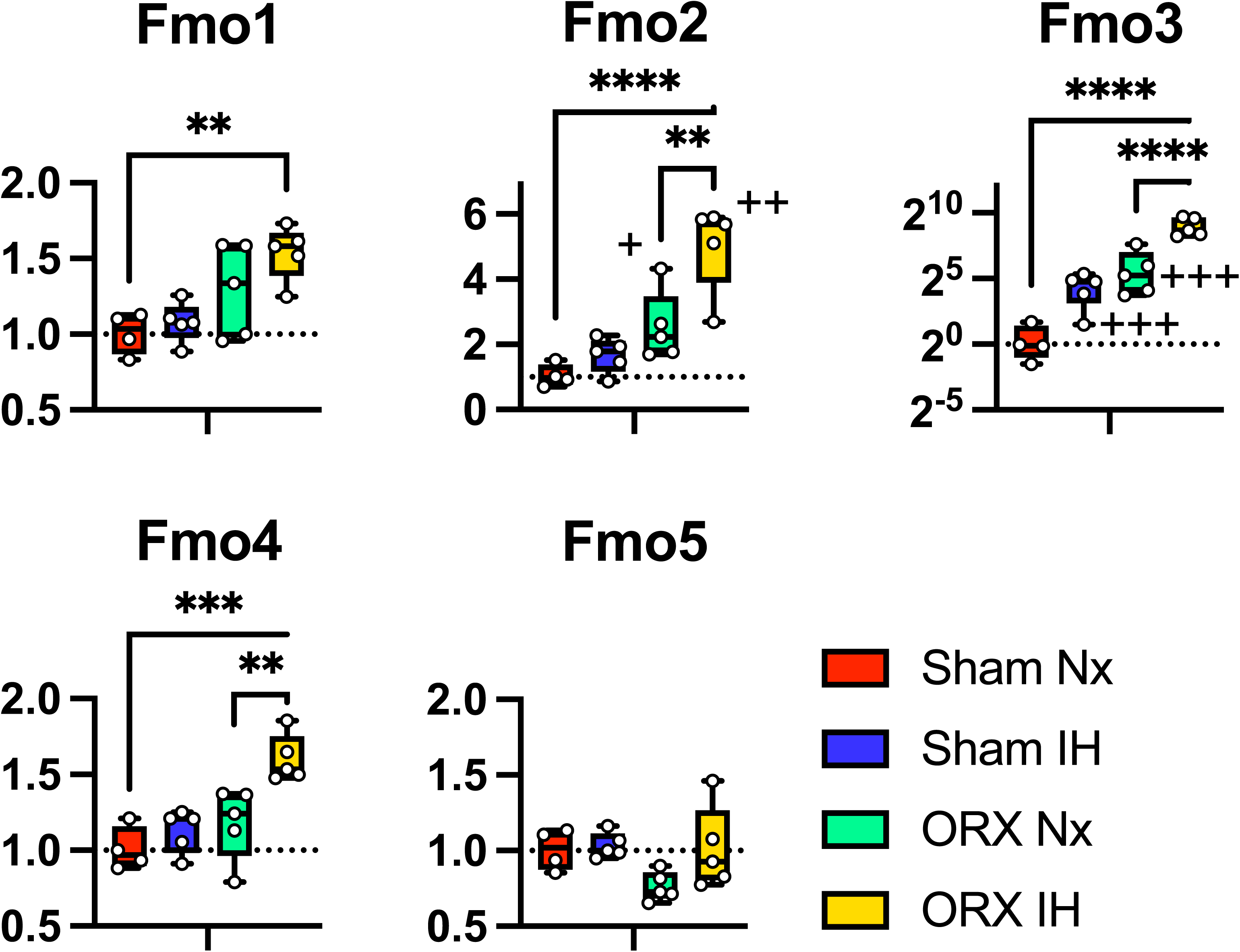
Relative expression of FMO 1-5 in the liver of Sham and ORX mice exposed to Nx or IH. All values are fold change relative to Sham Nx. P-values for post-hoc analysis: **, ***, ****: p<0.01, p<0.0001, and p<0.0001 IH vs Nx. +, ++, +++: p<0.5, p<0.01, and p<0.001 ORX vs Sham.

**Table 2:** Normalized counts of RNA reads (Trimmed mean of M-values – TMM) of Flavin-containing monooxygenase (FMO) 1-5, 6 and 9 in Sham and ORX mice exposed to Nx or IH. All values are mean ± SD. P value for post-hoc analysis: *, **, ****: p < 0.05, < 0.01 and <0.0001 IH vs Nx / †, ††, ††††: p < 0.05, < 0.01, and <0.0001 ORX vs Sham / $$, $$$, $$$$: p<0.01, <0.001 and < 0.0001 ORX IH vs Sham Nx.

|  | Sham Nx | Sham IH | ORX Nx | ORX IH |
| --- | --- | --- | --- | --- |
| FMO1 | 7.20 ± 0.20 | 7.30 ± 0.18 | 7.54 ± 0.35 | 7.82 ± 0.18<br>\$\$ |
| FMO2 | 2.83 ± 0.46 | 3.52 ± 0.54 | 4.11 ± 0.55<br>† | 5.13 ± 0.48<br>*, ††, \$\$\$\$ |
| FMO3 | -1.69 ± 1.31 | 2.38 ± 1.53<br>**** | 3.65 ± 1.56<br>†††† | 7.24 ± 0.68<br>****, ††††, \$\$\$ |
| FMO4 | 1.71 ± 0.20 | 1.87 ± 0.19 | 1.92 ± 0.33 | 2.38 ± 0.13<br>\$\$ |
| FMO5 | 7.58 ± 0.20 | 7.63 ± 0.11 | 7.18 ± 0.18 | 7.57 ± 0.36 |
| FMO6 | -7.39 ± 0.13 | -5.89 ± 1.19<br>** | -6.08 ± 1.57<br>†† | -5.24 ± 1.11<br>\$\$\$\$ |
| FMO9 | -8.07 ± 0.13 | -7.76 ± 0.46 | -8.09 ± 0.38 | -8.04 ± 0.48 |

## DISCUSSION

This study demonstrates that there are profound interactions between testosterone and exposure to intermittent hypoxia: on 1. the regulation of hepatic oxidative stress and glutathione metabolism at the level of enzyme activity; and 2. the regulation of hepatic metabolism, oxidoreductase activities, cytochromes dependent pathways, and glutathione metabolism at the level of gene expression. In mice exposed to IH the decreased body weight likely contributes to alter hepatic fat metabolism and induce oxidative stress, ultimately leading to the development of liver diseases (Gaucher et al., 2022; J. Huang et al., 2023). Our data show that the combined exposures to IH and ORX induce a unique enzymatic pro-oxidant profile with higher activity of the prooxidant enzyme NADPH oxidase, and reduced SOD activity. However, we have not been able to detect increased oxidative stress markers in ORX-IH mice compared to other groups, either by measuring the levels of oxidized lipids (MDA) or the total glutathione pool. To better understand this striking discrepancy, we applied a non-biased approach by assessing the transcriptomic profile of hepatic gene expression using RNA sequencing and pathway enrichment analysis. We identified significant enrichment for oxidoreductase activities, cytochromes dependent pathways, and glutathione metabolism. Of particular interest, one of the genes that showed the strongest up-regulation in ORX-IH mice is FMO3, a monooxygenase that has recently been shown as being critical to avoid oxidative damage in hepatic cells (S. Huang et al., 2021). Interestingly the cytochrome P450 3a44 (Cyp3a44) displayed a similar pattern of expression, and it is noteworthy that FMO and cytochromes are both acting as oxygenase and contribute to liver metabolism.

Previous studies in mice have shown that IH increases the levels of superoxide radicals and hydrogen peroxide and induces DNA breaks damage in the liver (Gaucher et al., 2022), which are clear signs of oxidative stress. These damages are accompanied by the activation of the classical antioxidant response pathways that includes higher expression of NRF1 and NRF2. The activation of the NRF1 pathway leads to a reconfiguration of the mitochondrial complexes that increases mitochondrial oxidation of fatty acids, while the activation of the NRF2 pathway increases the expression of SOD1 (cytosolic) and SOD2 (mitochondrial) (Gaucher et al., 2022), thereby increasing antioxidant defense. While our results do not show enhanced oxidative stress damage and the transcriptomic analysis do not show specific enrichment for these classical antioxidant defense pathways, cytosolic SOD and catalase activities are higher in Sham mice exposed to IH compared to their normoxic controls, strongly suggesting efficient antioxidant defense mechanism. One of the reasons explaining such discrepancy between the present transcriptomic analyses and the previous study (Gaucher et al., 2022) might be that we used a moderate frequency of exposure to IH (12 cycles/h in our study, vs 60 cycles/h in Gaucher et al., 2022). Another potential explanation is that, in the present study, mice were sacrificed several hours after the last IH cycles, therefore we assessed long-term rather than immediate responses. Nonetheless, the present results emphasize the temporal robustness of cytosolic SOD activation at the functional level of enzyme activity after IH exposures.

Contrasting with the results obtained in Sham mice, in ORX mice the activities of cytosolic and mitochondrial SOD were reduced by IH and the cytosolic NOX/SOD and NOX/CAT activity ratio were elevated. These results suggest that orchiectomy abrogates the responses that help prevent oxidative stress during IH exposures. This is consistent with studies showing that in rats testosterone supplementation reverses hepatic oxidative stress induced by aging, increases the activity of the antioxidant enzymes GPx, CAT and SOD, and prevents liver fibrosis (Zhang et al., 2019). This is also consistent with other data showing that testosterone increases the activity of antioxidant enzymes and reduces MDA in the blood and heart of diabetic rats (Chodari et al., 2019). Yet, and despite this apparent failure of antioxidant defense mechanisms in ORX-IH mice, there are no signs of hepatic oxidative damage. However, we believe that the results of the transcriptomic analysis provide the elements of a hypothetical framework involving the overexpression of flavine-containing monooxygenases (FMO) as an alternative antioxidant defense mechanism in ORX-IH mice.

Our data show that the expression levels of FMO 1-4 are elevated in the liver of ORX mice exposed to IH, showing synergistic effects between IH and ORX to regulate the expression of these enzymes. Negative regulations of FMO3 expression (both protein and mRNA) by testosterone in the liver of mice have been previously reported (Falls et al., 1997). To our knowledge, however, this is the first description of enhanced expression of hepatic FMO3 induced by IH and of a synergistic effect between ORX and another condition such as IH for the expression of FMO 1-4. FMO is a family of enzymes that are widely conserved from bacteria to mammals and are well known to facilitate elimination of oxidative xenobiotics, drugs, or endogenous molecules (Rendić et al., 2022). In this classical scheme of functions, these enzymes act by adding an oxygen molecule to nitrogen or sulfur atoms, thereby increasing the solubility of their substrates. The FMO family shares this detoxification function with cytochromes P450, and both families also share similar tissue and cellular localization as well as partial substrate specificity (Hao et al., 2009). In humans, FMO3 is essential to convert trimethylamine, a volatile substrate derived from the gut microbiota to the soluble trimethylamine-N-Oxide, and mutations in the FMO3 gene is associated with trimethylaminuria, a rare metabolic disorder (Phillips & Shephard, 2008). Even though the oxygenation reaction catalyzed by FMO can generate superoxide radical and hydrogen peroxide (Catucci et al., 2019; Siddens et al., 2014), recent studies in vertebrate and invertebrate species have shown that the most prevalent FMO enzymes (FMO 1-5) mediate beneficial effects by regulating metabolism and oxidative stress. In *c-elegans*, *fmo-2* mediates the increased lifespan induced by hypoxia and dietary restriction (Leiser et al., 2015). More recently, it is has been established that the *c-elegans fmo-2* gene induces a metabolic reconfiguration involving tryptophane as a direct substrate, and one carbon metabolism as a contributing pathway to prolong lifespan (Choi et al., 2023). In mice, caloric restriction markedly increases mRNA and protein hepatic FMO3 expression, and FMO3 overexpression mimics the robust beneficial antioxidant effects of caloric restriction on the liver, including a decreased MDA level and increased SOD activity (Guo et al., 2020). In vitro models have also shown that expression of FMO 1-5 improves survival of hepatic cells in response to several stressors, including the potent oxidant drug paraquat and the mitochondrial inhibitor rotenone (S. Huang et al., 2021). Interestingly, and despite this potent antioxidant functions of FMO and their increased expression in oxidative conditions, it has been reported that the expression of FMO is not regulated by NRF2 (Rudraiah et al., 2014). This is consistent with our own results, since there was no specific enrichment of pathways regulated by NRF2 in the RNAseq analysis. Overall, these results indicate that FMO 1-5 are important factors that regulate hepatic metabolism and prevent damage induced by a large variety of oxidative molecules. With this extensive background on the potent antioxidant effect of FMO enzymes, it is tempting to speculate that their increased expression in the liver of ORX-IH mice helps prevent MDA accumulation despite a prooxidant profile of enzyme activity.

## CONCLUSION

This study suggests that low levels of testosterone in male mice determine the mechanisms by which the liver handles IH-induced oxidative stress by increasing the expression of FMO enzymes to counterbalance a decreased SOD activity. This should be formally tested, but if future experiments prove that this is indeed the case this might likely open new avenues to better understand the consequences of IH in sleep apnea patients in terms of oxidative stress and control of redox status. As previously reported (Falls et al., 1997), our data show that the expression of FMO3 is regulated by testosterone, while it has been reported that estradiol also regulates its expression (Esposito et al., 2014), and that there are regulations during postnatal development (Janmohamed et al., 2004). Accordingly, and because sleep apneas have sex-specific consequences (Heinzer et al., 2018) and frequently occur in preterm neonates (Di Fiore et al., 2013) and children (Marcus et al., 2012), it would be necessary to explore the sex and age specific regulation of FMO expression and functions under IH. This could contribute to better understand protective mechanisms against the deleterious consequences of sleep apnea and help reduce the associated burden.

## Acknowledgments

Study funded by the Canadian Institutes of Health Research (Funding Reference Number: 162232). AB, GGC and VJ are members of the Quebec Respiratory Health Network.

## Declarations of interest

none

## Notes

### Competing Interest Statement

The authors have declared no competing interest.

